# An S-cone circuit for edge detection in the primate retina

**DOI:** 10.1101/667204

**Authors:** Sara S. Patterson, James A. Kuchenbecker, James R. Anderson, Andrea S. Bordt, David W. Marshak, Maureen Neitz, Jay Neitz

## Abstract

Midget retinal ganglion cells (RGCs) are the most common RGC type in the primate retina. Their responses mediate both color and spatial vision, yet the specific links between midget RGC responses and visual perception are unclear. Previous research on the dual roles of midget RGCs has focused on those comparing long (L) vs. middle (M) wavelength sensitive cones. However, there is evidence for several other rare midget RGC subtypes receiving S-cone input, but their role in color and spatial vision is uncertain. Here, we confirm the existence of the single S-cone center OFF midget RGC circuit in the central retina of macaque monkey both structurally and functionally, by combining single cell electrophysiology with 3D electron microscopy reconstructions of the upstream circuitry. Like the well-studied L vs. M midget RGCs, the S-OFF midget RGCs have a center-surround receptive field consistent with a role in spatial vision. While spectral opponency in a primate RGC is classically assumed to contribute to hue perception, a role supporting edge detection is more consistent with the S-OFF midget RGC receptive field structure and studies of hue perception.

## Introduction

Anatomical evidence for S-OFF midget RGCs is seen in the macaque (Old World) monkeys^38, 42, 72^ (but see Kolb et al.^40^) but not in New World monkeys.^45, 46^ Multi-electrode recordings in the macaque far peripheral retina find weak S-cone input to OFF midget RGCs, consistent with non-selective input from S-OFF and L/M-OFF midget bipolar cells;^32^ however, pure S-cone center, OFF midget RGCs have remained elusive in single cell electrophysiology^20, 48, 69^ and electroretinography.^43, 44^ The discrepancies in evidence for S-OFF midget RGCs may be due to differences in species,^52^ retinal eccentricity and methodology, thus, here, we sought both anatomical and physiological confirmation in the macaque retina. Because color vision,^53^ spatial acuity^71^ and midget RGC response properties vary considerably with eccentricity,^65^ we focused our efforts on the central retina where our research is most relevant to human visual perception.

The existence and function of S-OFF midget RGCs is relevant to a major unsolved question: how and where color and spatial information are separated in the visual pathway. This has been proposed to be accomplished in the cortex by theoretical downstream “de-multiplexing” circuits^24, 26^ (but see Kingdom & Mullen^37^) or in the retina by separate populations of RGCs for color and spatial information.^10, 61, 63^ In the central retina, each L- and M-cone provides the sole direct input to an ON and OFF midget circuit, forming a “private line” pathway from single cones to the parvocellular lateral geniculate nucleus (LGN).^41^ Their center-surround receptive fields perform a simple computation: comparing a single L-or M-cone center to neighboring L/M-cones in the surround. However, this seemingly simple computation introduces a complication for neural coding that vision science has yet to resolve. The center and surround receptive fields differ in both spatial location and spectral sensitivity, creating a neuron with both spatial and spectral opponency. The result is that L vs. M midget RGCs carry both spatial and spectral information, but confound the two such that from an individual midget RGC’s spike output, downstream neurons cannot distinguish between chromatic and spatial stimuli (see in Patterson et al., in revision). The goal of this work is not only to confirm the existence of S-OFF midget RGCs but understand the details of their circuity and receptive field properties so they can be fit into a larger understanding of the function of midget RGCs in color and spatial vision.

## Results

### S-cone circuits of the outer retina

To confirm the existence of S-OFF midget RGCs using serial electron microscopy (EM; Figure 1A), a first step was to reliably identify S-cones. We identified candidate S-cones by their small size and lack of long telendondria, branches forming gap junctions with neighboring cones.^40^ While S-cones were indeed smaller than L/M-cones, distinguishing the two at any single section was difficult, especially given the changing landscape of the foveal slope in our sample (Figure 1B, 1C, 1D).

**Figure 1.**
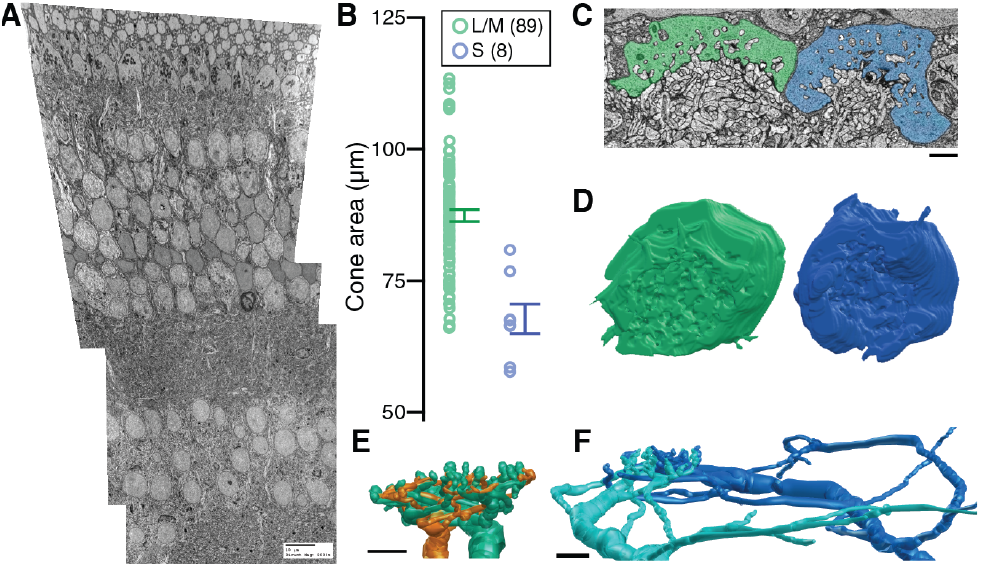
Identification of L/M- and S-cones using serial EM. **(A)** Transmission EM image of the block of tissue (magnification 2000x). **(B)** Area of L/M- and S-cone pedicles (S: 67.705 µm^2^ ± 2.81, LM: 87.359 µm^2^ ± 1.126; p = 0.0013). **(C)** Electron micrograph of neighboring LM-(green) and S-cones (blue). Scale bar is 2 µm **(D)** 3D reconstructions of neighboring S- and L/M-cones (blue, green). **(E)** 3D reconstruction of L/M-ON (teal) and L/M-OFF (orange) midget bipolar cell dendrites at an L/M-cone. **(F)** 3D reconstruction of S-ON bipolar cell dendrites at an S-cone.

Candidate S-cones were verified by reconstructing post-synaptic neurons. Cones signal changes in photon catch by modulating the rate of glutamate release from ribbon synapses onto a post-synaptic “triad” consisting of an ON-bipolar cell and two horizontal cells (reviewed by Sterling & Matthews^67^). L/M- and S-cones contact stereotyped horizontal cell and ON bipolar cell subtypes, allowing unambiguous confirmation of cone type by reconstructing the outer retina circuitry.

In the central retina, L/M-cones are densely innervated by a single ON midget bipolar cell while S-cones provide input to several S-ON bipolar cells (Figure 1E,1F).^42, 72, 76^ Furthermore, S-ON bipolar cells contact multiple S-cones, forming the distinctive lateral branches shown in Figure 2C. We reconstructed 14 S-ON bipolar cells (Figure 2A), each contacting up to three S-cones but receiving the majority of their input from a single S-cone.

**Figure 2.**
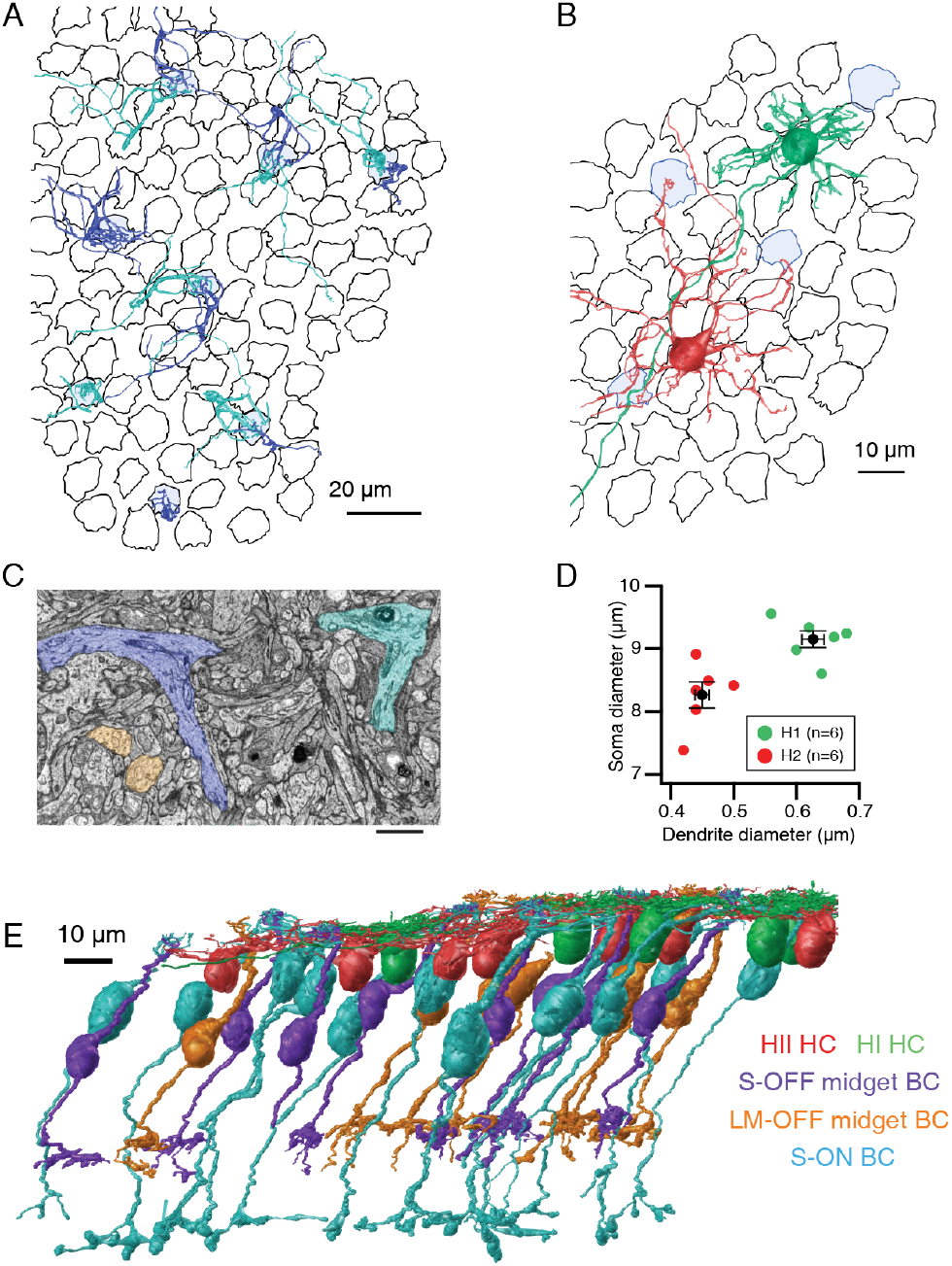
Serial EM reconstruction of the S-cone circuitry in macaque central retina. (A) 3D reconstructions of the dendrites of 14 S-ON bipolar cells over the cone mosaic (S-cones in blue). (B) 3D reconstructions demonstrate the morphological differences between HI (green) and HII (red) horizontal cells (S-cones in blue; L/M-cones in black). (C) Two S-ON bipolar cell dendrites (blue, cyan) converge towards an S-cone. These lateral branches distinguish S-ON bipolar cells from ON midget bipolar cells (orange) which branch directly beneath the L/M-cone pedicle. Scale bar is 2 µm. (D) Soma diameter plotted against primary dendrite diameter for HI and HII horizontal cells (n=6). The dendrite diameters are 0.627 µm ± 0.018 and 0.450 µm ± 0.011 for HI and HII horizontal cells, respectively (p ≤ 0.001). Soma diameters were 9.148 µm ± 0.134 and 8.259 µm ± 0.211 for HI and HII horizontal cells, respectively. (E) 3D reconstructions of the retinal neurons in this study.

The primate retina contains two horizontal cell subtypes: HII horizontal cells preferentially contact S-cones while HI horizontal cells avoid S-cones entirely.^1, 11, 33^ We reconstructed both types (Figure 2B) and confirmed each S-cone was densely innervated by HII, but not HI horizontal cells. In past light microscopy experiments, the horizontal cell subtypes could be distinguished by their dendritic field size and cone contacts, however, applying these features to serial EM experiments requires considerable annotation efforts. To date, no complete serial EM reconstructions of primate horizontal cells have been published. We developed two criteria for early identification of horizontal cell subtypes: soma size and primary dendrite diameter (Figure 2D). The horizontal cells identified using these criteria matched established morphological descriptions from light microscopy.^5, 39^

### The parvocellular LGN receives input from single S-cones through an S-OFF midget circuit

Having identified eight S-cones by morphology and postsynaptic circuitry, we next searched for OFF midget bipolar cell contacts. Indeed, a single OFF midget bipolar cell contacted each S-cone, as shown in Figure 3B. To verify the transmission pathway from S-cones to OFF midget RGCs, S-OFF bipolar cell contacts to S-cones were characterized. We next verified that S-OFF bipolar cells provide the sole input to OFF midget RGCs and confirmed the S-cone signal is not diluted by L/M-cone input from multiple midget bipolar cells, as seen in the peripheral retina.^32, 73^ Six OFF midget bipolar cells contacting neighboring L/M-cones were used as controls.

**Figure 3.**
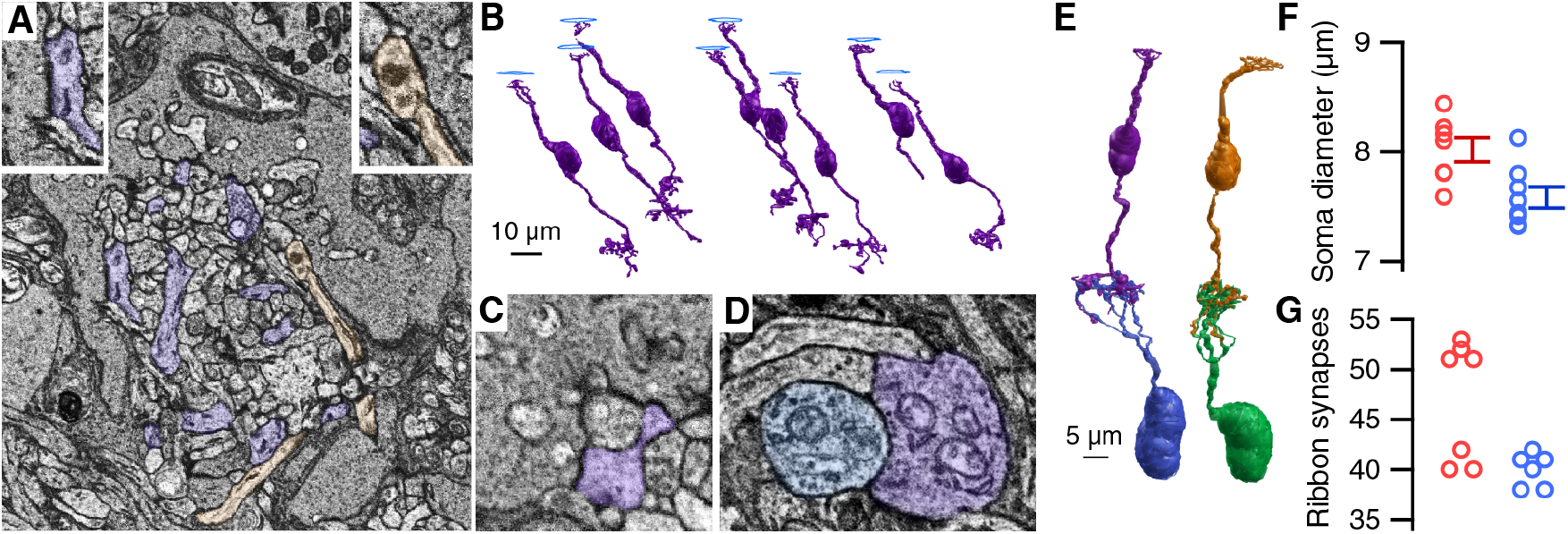
Each S-cone in the central retina provides the sole input to an OFF-midget circuit. **(A)** S-OFF midget (purple) and OFF diffuse (orange) bipolar cell processes at an S-cone. Left inset: An S-OFF midget bipolar cell basal synapse. Right inset: an OFF diffuse bipolar cell basal synapse. **(B)** 3D reconstructions of the S-OFF midget bipolar cells contacting eight S-cones. **(C)** An S-OFF midget bipolar cell dendrite at the triad-associated position. **(D)** An S-OFF midget bipolar cell making ribbon synapses onto an OFF midget RGC dendrite in the IPL. **(E)** Comparison of an S-cone OFF midget bipolar cell circuit (left) with an L/M-cone OFF midget circuit (right). **(F)** Soma diameters for S- and LM-cone OFF midget bipolar cells (S: 8:02 µm ± 0.11, n=7; LM: 7.93 µm ± 0.36, n=9, p=0.0175). **(G)** Ribbon synapses between OFF-midget bipolar cells and OFF-midget RGCs. S-OFF midget bipolar cells made 40 ± 0.683 ribbon synapses. LM-OFF midget bipolar cells formed two groups of 39.67 ± 0.577 (n = 3) and 51.75 ± 1.291 (n = 4) ribbon synapses (expressed as Mean ± SD). The probability obtaining these two groups by chance from a single normally-distributed group was estimated using a bootstrap procedure (p=0.0083, see Methods).

OFF bipolar cells for “flat” basal synapses along the base of the cone pedicle. Basal synapses are recognized as membrane densities without associated pre-synaptic vesicles as OFF bipolar cells respond to glutamate released by the ribbon synapses. Synaptic contact was quantified at two S-cones, finding 22 and 23 basal synapses. The basal synapse counts are likely underestimates as the 90 nm section thickness prevented exhaustive tracing of all bipolar cell dendrites through the S-cone pedicle. However, these numbers are comparable to Klug et al. (2003), and to the 25 OFF midget bipolar cell basal synapses counted at a neighboring L/M-cone.

Each OFF bipolar cell subtype is located at a characteristic distance from the ribbon synapse (Figure 3C). At L/M-cones, OFF midget bipolar cell contacts are located at the “triad associated” position, adjacent to the membrane invaginations containing horizontal cell and ON bipolar cell dendrites.^9, 35, 72^ Virtually all S-OFF midget bipolar cell synapses were found in the same location, adjacent to the membrane invaginations containing S-ON bipolar cell dendrites (Figure 3C). This distinguished the S-OFF midget bipolar cell dendrites from the thin, straight, OFF diffuse bipolar cell dendrites making basal synapses around the edges of the S-cone pedicle, as in Figure 3C. The distance from the ribbon synapse shapes bipolar cell responses by defining the timing and concentration of available glutamate.^28, 58^ Thus S-OFF midget bipolar cells likely have similar response properties to L/M-OFF midget bipolar cells.

Near the fovea, each midget bipolar cell contacts a single midget RGC, forming a “private line” from a single cone to the parvocellular LGN. Outside the central retina, the number of bipolar cells converging on a single midget RGC scales with eccentricity.^16, 41^ We traced seven of the S-OFF midget bipolar cells to the inner retina, where each contacted a single OFF midget RGC with an average of 40 ribbon synapses (Figure 3D). The eighth S-OFF midget bipolar cell ran off the edge of the volume. Ribbon synapses from S-OFF midget bipolar cells onto other RGCs were rarely observed.

Like the S-OFF midget bipolar cells, each L/M-OFF midget bipolar cell provided the sole input to an OFF midget RGC. Ribbon synapse counts have been reported to divide L/M-OFF midget bipolar cells into two populations, referred to as “sparsely” and “densely” branching, which presumably correspond to L- and M-cones^7, 8^ (but see Schein et al.^62^). The L/M-OFF midget bipolar cells in our study made an average of 39 or 51 ribbon synapses, with the sparsely branching group comparable to the S-OFF midget bipolar cells (Figure 3G). The only anatomical difference was that S-OFF midget bipolar cells had slightly smaller somas than the LM-OFF midget bipolar cells (Figure 3F).

Overall, we found no anatomical evidence for a significant functional difference between L/M-OFF and S-OFF midget circuits. Taken together, our reconstructions indicate the parvocellular LGN indeed receives input from single S-cones through S-OFF midget RGCs in the central retina.

### The parvocellular LGN receives input from single S-cones through an S-OFF midget circuit

Previous studies may have missed rare midget RGCs with S-cone input due to the unique technical challenges involved in studying S-cone circuitry. Short-wavelength light is attenuated by almost everything in the optical light path: lenses, objectives and even the macular pigment of the retina itself. Furthermore, the typical 465 nm ‘blue’ LED in standard CRT displays stimulates the M-cones near as much as the S-cones. Control over the spectral distribution of a light source typically requires sacrificing the spatial resolution provided by commercial projectors, limiting experiments to full-field stimuli. To overcome these challenges, we created a custom light source specifically designed to minimize short-wavelength attenuation while maximizing S-cone contrast by replacing the built-in 465nm LED in a Lightcrafter DLP with a 405nm LED. Thus, we optimized S-cone isolating stimuli while maintaining the projector’s spatial resolution.

During the course of a larger series of electrophysiology experiments on S-cone inputs to midget RGCs using our custom-built light source, we encountered four midget RGCs with S-OFF responses and single cone receptive field centers (Figure 4A, 5C). A temporally-modulated S-cone isolating square-wave was used to assess S-cone inputs for every RGC encountered. The S-cone isolating stimulus modulates S-cone activity while holding L- and M-cones constant.^31^ As shown in Figure 4A, each potential S-OFF midget RGC responded robustly to the S-cone decrements.

**Figure 4.**
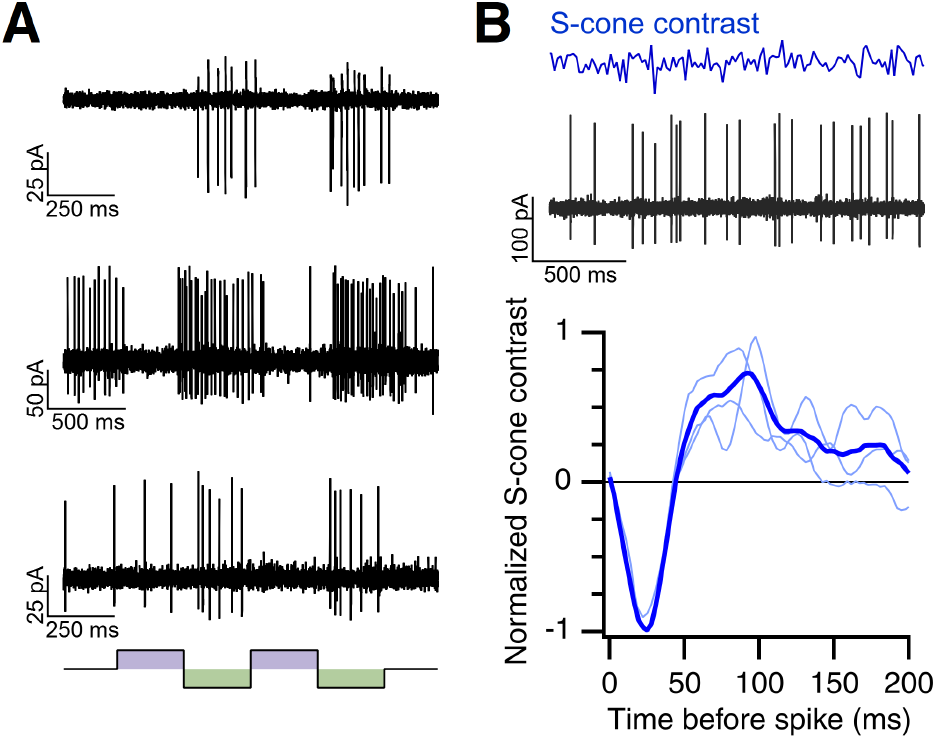
Spike responses of S-OFF midget RGCs. **(A)** Responses of three OFF midget RGCs to a temporally-modulated S-cone isolating spots. **(B)** An example of the S-cone isolating Gaussian white noise (top) and an S-OFF midget RGC’s response (middle). Bottom: The average linear filter (blue) from three S-OFF midget RGCs (light blue). The linear filter represents the average S-cone contrast preceding a spike.

The S-cone response kinetics were characterized using a time-varying "white noise" stimulus where the S-cone contrast of each frame was drawn pseudo-randomly from a Gaussian distribution (mean = 50%, SD = 30%; Figure 4B). Convolving the stimulus with the elicited spike rate histogram in the Fourier domain returns a ‘filter’ that captures the linear components of the response. In other words, the linear filter represents the average S-cone modulation preceding a spike and is proportional to the neuron’s impulse response function.^12, 59^ The average negative peak at 25 ms in Figure 4B indicates that the neuron fires following decrements in S-cone contrast, the defining characteristic of an “S-OFF” neuron.

Our anatomical data confirm that HII horizontal cells carry both L/M-and S-cone signals (Figure 2B).^18, 33^ While an HII horizontal cell mediated L/M-cone surround has been reported in individual S-cones,^55^ feedback from the strong S-cone surround has not been reported. Consistent with the presence of HII horizontal cell-mediated S-cone feedback, the S-OFF midget RGCs responded weakly to large, full-field S-cone isolating stimuli, preferring small spots centered over the receptive field (Figure 5A). The weak responses to full-field stimuli may help explain why previous electrophysiology studies did not find evidence of S-OFF midget RGCs.^69^

**Figure 5.**
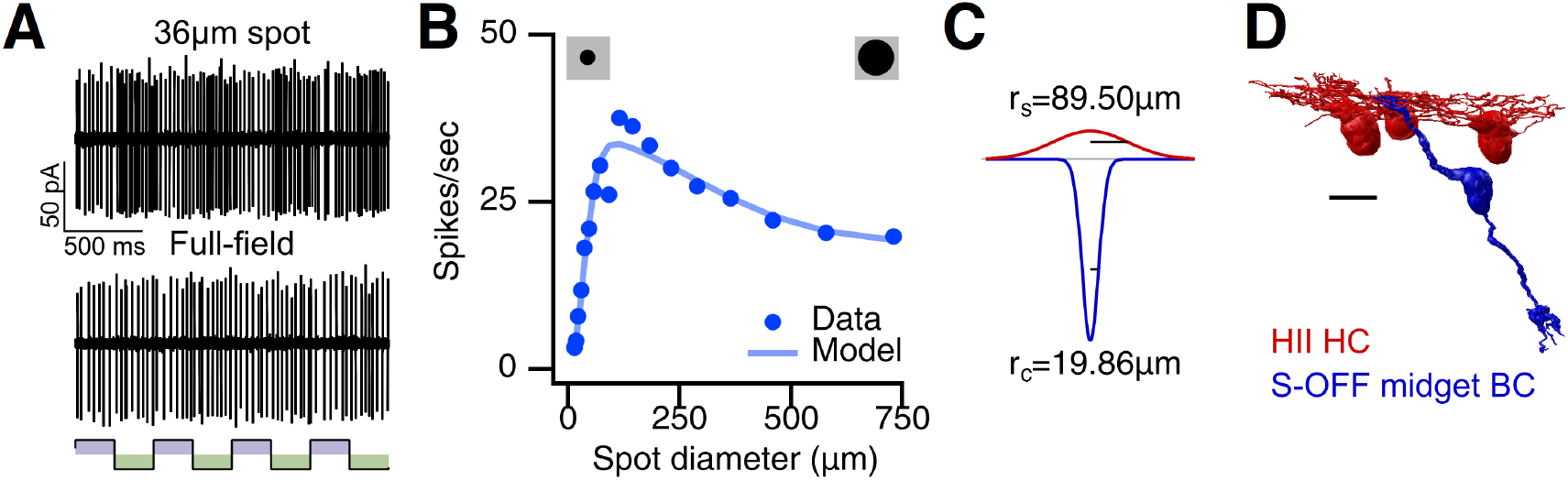
S-OFF midget RGCs encode spatial information with a center-surround receptive field. **(A)** Responses of a fourth OFF midget RGC to a 36 µm spot vs a full-field stimulus (both S-cone isolating square-wave at 2 Hz). **(B)** Responses to spots of increasing diameter presented as temporal modulations (4 Hz square-wave) achromatic spots of increasing diameter. Smooth curves are fits to a Difference of Gaussians model (Kc=8.28 spikes/sec, Rc=19.86 µm, Ks =5.82 spikes/sec, Rs=89.5 µm). **(C)** The S-OFF midget receptive field obtained by the Difference of Gaussians fit in **B**. **(D)** The anatomical basis for the S-cone center-surround receptive field. The center receptive field represents the single S-cone input directly to the S-OFF midget bipolar cell (blue). HII horizontal cell feedback (red) forms the antagonistic surround receptive field. Scale bar is 10 µm.

The center-surround receptive field structure was investigated using an expanding spot stimulus temporally modulated in luminance (Figure 5B). The spike rate increased as the spot size expanded to cover the entire center receptive field, then began decreasing as the spot further expanded to cover more of the antagonistic surround receptive field. The strength and extent of the center and surround receptive fields were determined by fitting the F1 amplitude to a standard Difference of Gaussians model (Figure 5C, Kc = 8.28 spikes/sec, Rc = 19.86 µm, Ks = 5.82 spikes/sec, Rs = 89.5 µm).^13, 14, 30^ The fit parameters compare well with measurements of the S-cone mosaic in our serial EM volume. The 19.86 µm center receptive field radius falls within the 33.83 ± 1.38 µm (n=8) nearest neighbor distance of the S-cone mosaic, indicating these do reflect the responses of single cone center midget RGCs. The tuning profile of spatial opponency clearly distinguished S-OFF midget RGCs from melanopsin RGCs, the other known S-OFF neuron in the primate retina.^19^ Taken together, the S-OFF midget RGC anatomy and physiology indicates a center-surround receptive field, similar to L/M-OFF midget RGCs (Figure 4B, 5A-D).

The relationship between center-surround receptive fields, spatial information and edge detection is well-studied.^36, 51, 56, 66^ Thus, the presence of center-surround structure in S-OFF midget RGCs indicates that, regardless of spectral tuning, the S-OFF midget RGC is carrying spatial information.

## Discussion

Our work provides the most comprehensive reconstruction of the primate outer retinal S-cone circuitry to date, including the first complete reconstructions of horizontal cells in the primate retina. To the best of our knowledge, this work is also the first single cell electrophysiology experiment to demonstrate an S-cone surround receptive field in a primate RGC, as would be predicted from HII horizontal cell circuitry. While the presence of an S-cone surround receptive field at individual S-cones is an expected consequence of the HII horizontal cell feedback, the resulting spatial opponency has not been tested nor incorporated into the most popular models of S-cone circuits.

Here, we find that S-OFF midget RGCs have the same center-surround receptive field structure as L vs. M midget RGCs. This receptive field structure is consistent with the edge detection required for high acuity spatial vision.^51^ It has been argued that edge detection must be achromatic and any degree of spectral opponency is detrimental.^3, 50^ However, not all edges are defined by changes in intensity alone, and equiluminant edges are common in natural scenes.^34^ Thus, the spectral opponency in S-OFF midget RGCs could be used to signal the presence of an edge defined by wavelength or intensity. At equiluminance, we need S-cones to detect white-yellow and gray-brown boundaries.^6^ Transitions from white to yellow and gray to brown both involve S-cone decrements so S-OFF midget RGCs are ideally suited for signaling brown or yellow objects against neutral backgrounds. Also, seeing light-colored objects against the blue sky may be mediated by S-OFF RGCs.

S-OFF midget RGCs have been proposed to mediate yellow hue percepts, forming the OFF counterpart to the small bistratified RGC,^17^ despite extreme asymmetries between the two types. However, because their center-surround receptive fields confound color and spatial information, hue cannot be extracted from individual S-OFF midget RGCs. The same problem exists for L/M midget RGCs, however, at least theoretically, the problem could be solved by combining the signals from L center/ M surround and M center/L surround midget RGC.^24, 26^ An analogous solution is not available for S-OFF midget RGCs because there are no L/M cone center S-cone surround midget RGCs. The simple conclusion is that S-OFF midget RGCs are specialized edge detectors used for spatial vision. They can detect edges based either on luminance or spectral contrast but are unlikely to have any role in hue perception. Moreover, the S vs. L+M spectral tuning in both neurons does not match the cone inputs to blue-yellow hue perception^23, 64, 68, 78, 79^ and a subset of midget RGCs with cone inputs matching the fundamental hues^21, 22, 27, 70^ has been proposed to mediate hue perception instead (for review, see Neitz & Neitz^54^). These L vs. M midget RGCs with significant S-cone input to the surround receptive field are another controversial midget RGC subtype that warrants additional investigation. The significant technical challenges involved in studying rare S-cone inputs to parafoveal midget RGCs that limited these investigations in the past can now be overcome with the multi-disciplinary approach described here.

## Methods

### Serial Electron Microscopy

#### Tissue Preparation

Retinal tissue was obtained from a terminally anesthetized macaque (*Macaca nemestrina*) monkey through the Tissue Distribution Program at the Washington National Primate Center. All procedures were approved by the Institutional Animal Care and Use Committee at the University of Washington. A 0.2 by 0.2 mm block of inferior parafoveal retinal tissue at ∼ 1 mm eccentricity from the fovea center were processed as previously described.^25^ At this eccentricity (the edge of the foveal slope), the displacement of RGCs from cone pedicles was minimized while still remaining in a region where most midget RGCs receive single cone input. A serial block-face scanning electron microscope (Zeiss/Gatan) was used to section and image the retinal tissue at a resolution of 7.5 nm/pixel. The volume contained 1893 90 µm sections from the ganglion cell layer through the cone pedicles. Image registration was performed using Nornir (http://nornir.github.io). The transmission EM image in Figure 1A was taken of the inferior retina volume prior to sectioning.

#### Annotation

The serial EM volumes were annotated using the web-based, multiuser Viking software described previously (http://connectomes.utah.edu).^2^ Briefly, processes were traced through the sections by placing a circular disc at the structure’s center of mass and linking the disc to annotations on neighboring sections. Cone pedicles were outlined using a closed curve polygon defined by three or more control points. Synapses were annotated with lines connected by 2-3 control points and linked to a parent neuron. Synapse identification used previously described parameters.^29, 72^

#### Data Analysis and Visualization

Data analysis and 3D rendering were performed using an open-source Matlab program (https://github.com/neitzlab/sbfsem-tools).^4^ The cone pedicle analyses were based on XYZ coordinates of the closed curve control points, connected by Catmull-Rom splines. All other analysis was performed using the X, Y, Z coordinates and radius of the Disc annotations. For Figure 1D, the closed curve coordinates were used to build a volume from which isosurfaces were extracted using the marching cubes algorithm and rendered as a triange mesh.^47^ All other 3D models are triangle meshes built by rendering segments of connected annotations as rotated cylinders centered at each annotations’ XYZ coordinates and scaled by their radii.

Soma diameter was calculated from the single largest annotation in each neuron, assumed to be the soma. Primary dendrite diameter was calculated as the median diameter of annotations centered 0.5-1.5 µm from the soma.

The probability that the reported distribution of ribbon synapses in Figure 3G was drawn by chance from a single normally-distributed group of L/M-cone OFF midget bipolar cell ribbon synapses was determined using a bootstrapping procedure. A normal distribution with the mean and standard deviation (SD) of both groups combined formed the null hypothesis. Seven integers were drawn from this normal distribution, then divided into two groups above and below the mean. The average SD of these two groups provided a metric for the degree of bimodality – two groups drawn from a normal distribution should have large SDs while two groups from two distinct distributions should have smaller SDs, as demonstrated by the clustering in Figure 3G. The percentage of 10,000 boostrap ribbon synapse distributions with equal or higher average SDs than the original dataset determined the reported p-value. All other reported statistics used the Mann-Whitney-Wilcoxon ranked sum.

### Electrophysiology

#### Tissue Preparation

Retinal tissue was obtained from terminally anesthetized macaque monkeys (*M. nemestrina*, *M. fasicularis*, *M. mulatta* of both sexes) through the Tissue Distribution Program at the Washington National Primate Center. All procedures were approved by the Institutional Animal Care and Use Committee at the University of Washington. Dissections were performed as previously described.^57^ Briefly, enucleated eyes were hemisected and the vitreous humor was removed mechanically. When necessary, the eye cup was treated for ∼ 15 minutes with human plasmin (∼ 50 µg mL^*−*1^, Sigma or Haematologic Technologies) to aid vitreous removal.

#### Recording

A piece of macaque macular retina with well-attached retinal pigment epithelium was placed on the stage of a microscope ganglion cell side up. The tissue was superfused with warmed (32-35°C) Ames’ medium (Sigma) at ∼6-8 mL min^*−*1^. In some cases, additional D-glucose (14 mmol) was added to the Ames’ medium.^49^ Ganglion cell spikes were measured with extracellular or loose-patch recordings using an Ames-filled borosilicate pipette. The data was sampled at 10 kHz (Multiclamp 700B, Molecular Devices), Bessel filtered at 3 kHz and digitized using an ITC-18 analog-digital board (HEKA Instruments).

#### Cell Identification and Selection

RGCs were initially identified by soma appearance, as visualized with a 60x objective (Olympus) under infrared illumination. RGC type was further determined by responses to spots, cone-isolating stimuli and mapping the receptive field with horizontal and vertical bars. Midget RGCs make up over 90% of all RGCs in the central retina^77^ and were confirmed by small soma, sustained responses and small center-surround receptive fields.^15, 80^ OFF RGC somas were generally vitread to ON RGC somas. In addition to these criteria, S-OFF midget RGCs were identified using small S-cone isolating stimuli positioned over the receptive field center.

#### Stimuli

Stimulus presentation and data acquisition used the open source programs Stage (www.stage-vss.github.io) and Symphony (www.symphony-das.github.io), respectively. The Symphony stimulus protocols used in this study can be found at https://github.com/sarastokes/sara-package. Visual stimuli were projected onto the cone outer segments through a 10x objective (Olympus) using a Lightcrafter DLP 4500 (Texas Instruments) with a 60 Hz frame rate. To optimize S-cone isolation, the built-in LEDs were replaced with custom LEDs at 405, 535 and 630 nm.

Each LED was calibrated by measuring the spectral distribution with a spectroradiometer (Konica Minolta CS-2000) and the power with an optometer (UDT-300). A transformation matrix, *A*, relating the LED weights to the cone quantal catches was obtained by taking the outer product of the LED spectra (*R*(*λ*), *G*(*λ*) and *B*(*λ*), as only three of the four LEDs were used at a time) and the L-, M- and S-cone spectral sensitivities (*L*(*λ*), *M* (*λ*), *S*(*λ*) respectively):

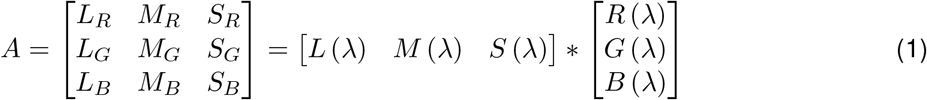

The transformation matrix was used to solve for the appropriate LED weights for any given level of L-, M- and S-cone activations.^31^

The mean light levels were calculated using a collecting area of 0.37 µm^2^ and 1 µm^2^ 77. All stimuli used photopic light levels (∼3 × 10^3^ to 3 × 10^5^ R ∗ /rod/s. To maintain a constant state of light adaptation, the mean light level was displayed continuously between stimulus presentations. Contrast is expressed as Weber contrast.

#### Recording Protocol

Tissue sensitivity was assessed at the beginning of each experiment by ensuring ON parasol RGCs responded to a full-field, 5% contrast, 4 Hz temporally-modulated spot.^74^ ON parasol RGCs lack significant S-cone input and were also used to validate the S-cone isolating stimuli.^32^

For each subsequent RGC encountered, the polarity (ON, OFF or ON-OFF) and cell type were first determined by spots presenting high contrast luminance increments and decrements from a photopic mean light level. Cell type was confirmed by receptive field dimensions. The receptive field center was determined by vertical and horizontal bars, presented as 2-4 Hz squarewave temporal modulations. The center and surround receptive field radii were measured from Difference of Gaussian fits to expanding spots and annuli. In cases where the receptive field dimensions were unclear (either due to noise or a rare cell with an atypical receptive field), the measurements were confirmed by estimating the spatiotemporal receptive field using coarse (25-50 µm square pixels) binary spatial noise.

S-cone contributions were measured in each RGC with different sizes of S-cone isolating spots, presented as 1-2 Hz squarewave temporal modulations. For each RGC with a significant S-cone response, the temporal and spatial characteristics were next measured with S-cone isolating full-field temporally modulated Gaussian noise and expanding spots, respectively.

##### Expanding Spot

The spot stimulus was positioned at the receptive field center and presented as a 4 Hz temporal modulation. Given the low contrast of the S-cone isolating stimulus, the spot stimulus was repeated as a high contrast luminance modulation.

##### Gaussian noise

The temporal ‘white noise’ stimulus was generated by psuedo-random draws from a Gaussian distribution centered at the mean light level (mean, 50%, SD = 30% contrast). The noise stimulus was presented in 10 or 20 second epochs, with 1 second interval between epochs.

#### Data Analysis

##### Difference of Gaussians model

A Difference of Gaussians (DoG) model was fit to the F1 amplitudes.^30, 60^ The DoG model characterizes the center and surround receptive fields as two antagonistic, two-dimensional Gaussians with separate strengths and sizes. The center and surround receptive fields are assumed to be radially symmetric and centered at the same location. The DoG model predicts the response, *R*, for spot diameter, *f* as:

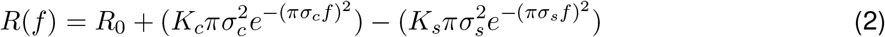

where *R*_0_ is the baseline response, *σ*_*c*_ and *σ*_*s*_ are the center and surround receptive field sizes, respectively, and *k*_*c*_ and *k*_*s*_ are the center and surround strengths, respectively.

##### White noise analysis

Analysis was performed on the spike rate, binned at 360 Hz for temporal noise and 120 Hz for spatiotemporal noise. The first second of the response was omitted to control for adaptation. The linear filter, *F*, is obtained by cross-correlating the stimulus, *s*(*t*), with the spike rate, *r*(*t*), and dividing out the stimulus’ power spectrum:

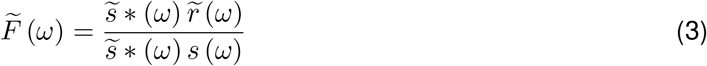

where 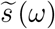 is the Fourier transform of, *s*(*t*), 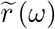 is the Fourier transform of *s*(*t*) and ∗ denotes the complex conjugate. In practice, the stimulus power spectrum was nearly flat and the denominator was omitted.^75^ The inverse transform of 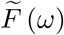 returned the time-domain linear filter, *F*.

## Acknowledgments

This work was supported by NIH grants EY027859 (J.N.), NS099578 (S.S.P.), EY07031 (S.S.P.), EY02576 (D.W.M.), EY015128 (J.R.A), EY028927 (J.R.A.), EY014800 (J.R.A.), EY001730 (Core grant for vision research). We thank Mark Cafaro, Shellee Cunnington, Toni Haun, Ed Parker and Jessica Rowlan for excellent technical support. Tissue was provided by the Tissue Distribution Program at the Washington National Primate Research Center (WaNPRC) with the help of Chris English. William Grimes, Michael Manookin, Fred Rieke, Raunak Sinha and Max Turner assisted with tissue preparation.

## Author Contributions

J.N. and S.S.P, conceived the experiments and wrote the initial draft of the manuscript. S.S.P, J.N., J.A.K., J.R.A., A.S.B., D.W.M. and M.N. contributed to the serial electron microscopy experiments and data analysis. S.S.P. performed the electrophysiology experiments. J.A.K., J.R.A., A.S.B., D.W.M. and M.N. read and contributed to the final version of the manuscript.

## Data Availability

Access to the macaque EM volume dataset is available on request. Visualizing both the dataset and the annotations requires the Viking Viewer developed in Bryan Jones’ lab at (http://connectomes.utah.edu). The 3D reconstructions from Viking Viewer annotations are visualized with SBFSEM-tools, an open-source Matlab toolbox developed in the Neitz lab (https://github.com/neitzlab/sbfsem-tools). Access to the dataset through the Viking Viewer and SBFSEM-tools does not require that the user install the data locally. The data and code used to generate each figure will be made available upon publication at https://github.com/neitzlab/SConeEdgeDetection.

